# Mental Simulation of Facial Expressions: Mu Suppression to the Viewing of Dynamic Neutral Face Videos

**DOI:** 10.1101/457846

**Authors:** Ozge Karakale, Matthew R. Moore, Ian J. Kirk

## Abstract

The mirror neuron network (MNN) has been proposed as a neural substrate of action understanding. Electroencephalography (EEG) mu suppression has commonly been studied as an index of MNN activity during execution and observation of hand and finger movements. However, in order to establish its role in higher order processes, such as recognising and sharing emotions, more research using social emotional stimuli is needed. The current study aims to contribute to our understanding of the sensitivity of mu suppression to facial expressions. Modulation of the mu and occipital alpha (8 - 13 Hz) rhythms was calculated in 22 participants while they observed dynamic video stimuli, including emotional (happy and sad) and neutral (mouth opening) facial expressions, and non-biological stimulus (kaleidoscope pattern). Across the four types of stimuli, only the neutral face was associated with a significantly stronger mu suppression than the non-biological stimulus. Occipital alpha suppression was significantly greater in the non-biological stimulus than all the face conditions. Source estimation (sLORETA) analysis comparing the neural sources of mu/alpha modulation between neutral face and non-biological stimulus showed more suppression in the central regions, including the supplementary motor and somatosensory areas, than the more posterior regions. EEG and source estimation results may indicate that reduced availability of emotional information in the neutral face condition requires more sensorimotor engagement in deciphering emotion-related information than the full-blown happy or sad expressions that are more readily recognised.

## 1 Introduction

Nonverbal communication is a crucial component of human social behaviour, but its neural mechanisms are poorly understood. The ability to understand others’ mental states from their facial and bodily gestures allows us to respond effectively during social communication. Gallese and Goldman (1998) proposed a simulation theory of action understanding to account for the complexity of this process. Under this model, on observing an action, the observer subconsciously and automatically employs a specialised neural circuitry to simulate the action using their own motor system, in turn activating mental states associated with execution of the action, and providing insight into the mental state of the actor. The neural substrate of the simulation theory is proposed to be the *mirror neuron* (Gallese & Goldman, 1998).

Mirror neurons were first discovered in the motor areas of the monkey brain (di Pellegrino et al., 1992). They were observed to fire during both execution and observation of actions, such as grasping an object, putting it in mouth or breaking it. Moreover, the sensory modality by which the action was experienced did not seem to matter for a subset of these neurons: they were triggered by the sound of the action, even when the action was not seen (Kohler et al., 2002). The implication of these findings was that the mirror neurons could be coding the representations of the actions, allowing for recognising the movements involved in an action and inferring the intention behind the action. Evidence for a similar mirroring mechanism in the human brain has come from functional magnetic resonance imaging (fMRI) and positron emission tomography (PET) studies (Caspers et al., 2010; Molenberghs et al., 2012) as well as single neuron recordings during surgery in humans (Mukamel et al., 2010).

In addition to metabolic brain imaging and *in vivo* cellular studies, EEG studies have measured mu rhythm desynchronisation to infer mirroring activity. Mu rhythm, characterised by 8-13 Hz oscillations detected over the sensorimotor area, is mostly associated with the functions of the sensorimotor cortex (Niedermeyer, 2005). Increased mu rhythm power indicates physical inactivity and resting, with movement execution as well as observation leading to its suppression (Cochin et al., 1999; Cochin et al., 1998; Fecteau et al., 2004; Hari & Salmelin, 1997; Lepage & Theoret, 2006; Muthukumaraswamy et al., 2004). Due to the responsivity of the mu rhythm to action observation, it has been proposed to reflect mirror neuron activity related to viewing of biological action with or without object interaction, including finger movements (Babiloni et al., 1999; Cochin et al., 1999) and hand grip movements (Muthukumaraswamy et al., 2004), as well as hearing sounds that are linked to actions, such as piano melodies (Wu et al. 2016).

Since the initial discovery of mirror neurons, research has focused on their potential role in social cognitive processes that rely on an ability to understand actions and intentions, such as empathy. Similar activity in the brain regions observed in fMRI during execution and observation of facial expressions has been suggested to provide evidence for the existence of a single mechanism of action representation which allows people to empathise with others (Carr et al., 2003). In order to further the knowledge about the role of the MNN in social emotional information processing, a group of researchers used EEG mu suppression as a proxy of the MNN to investigate the network’s sensitivity to emotional information using body parts in painful and non-painful situations, and found greater mu suppression in the painful compared to non-painful conditions (Cheng et al., 2014; Hoenen et al., 2015; Yang et al., 2009). In contrast to findings which suggest a heightened sensitivity of the mu rhythm to emotional information, others found similar levels of mu modulation during gender discrimination and emotion recognition tasks which entailed viewing point-light displays of human figures’ walk (Perry, Troje et al., 2010). Facial expressions have been used as stimuli in EEG mu suppression research only in a handful of studies (Cooper et al., 2013; Moore & Franz, 2017; Moore et al., 2012; Rayson et al., 2016; Rayson et al., 2017). Further research using different types of facial movements depicting varying levels of emotional information as the visual stimuli is necessary to investigate the differential sensitivity of the sensorimotor cortex to emotion-related information processing.

It is crucial to note that findings from some mu suppression studies indicate that mu can easily be confounded with occipital alpha activity, yielding alpha suppression at the central electrodes that is not only similar while viewing biological and non-biological motion, but also more pronounced to biological than non-biological motion when the observed action depicts pain. As pointed out by Milston et al. (2013), most of the studies that have explored the relation between mu suppression and empathy have used stimuli eliciting pain only (e.g., Hoenen et al., 2015; Perry, Bentin et al., 2010; Yang et al., 2009). Researchers have highlighted that processes other than empathy, such as attention, may be at work while viewing painful stimuli due to their threatening nature (Hoenen et al., 2013) or salience (Perry, Bentin et al., 2010). It may still be difficult to disentangle mu from alpha in tasks using non-emotional biological motion. For example, Aleksandrov and Tugin (2012) did not find any systematic differences in mu suppression to the observation of hand movements, non-biological objects or mental counting. Similarly, Perry and Bentin (2010) observed that alpha suppression at the mu and the occipital areas were very similar to the observation of hand movements toward an object. A recent study conducted by Hobson and Bishop (2016) showed that different types of baseline used to measure mu suppression engage the attention system differently, thus directly impacting the degree of suppression recorded. They found that mu and occipital alpha modulation while viewing hand movements and kaleidoscope movements were consistent with the MNN activity only when the static video of the image that immediately preceded the dynamic video of the image was used as the baseline. Due to the posterior alpha confound associated with attentional processes, the baseline and the control conditions need to be chosen carefully.

The current study aims to contribute to our understanding of the simulation account by investigating the responsiveness of the sensorimotor cortex to emotional and non-emotional facial expressions. Our goal is to examine the differential sensitivity of the mu rhythm to different types of facial movements. To our knowledge, this is the first study to examine the sensitivity of mu rhythm, while controlling for occipital alpha activity, to dynamic neutral and emotional facial expressions not depicting pain. A within-trial baseline method was adopted as per Hobson and Bishop (2016): the 1100 ms static image epoch was used as the baseline for quantifying activity in the subsequent 2050 ms dynamic image epoch. It was hypothesized that mu suppression would be greater in the (1) happy, sad and neutral face conditions than the non-biological stimulus condition, and (2) happy and sad face conditions than the neutral face condition, without a corresponding difference in occipital alpha suppression.

## 2 Materials and methods

### 2.1 Participants

Twenty-five participants (16 female) between the ages of 19 and 36 (*M* = 26.5, *SD* = ±6) were recruited through flyers placed around the University of Auckland campus. Each participant was compensated with a $20 supermarket voucher. Prior to data collection, a pre-screening questionnaire was emailed to the volunteers to identify whether they met the criteria for participation. Exclusion criteria included self-reported major head injury, psychiatric diagnosis, psychoactive medication use, or sensorimotor problems. All of the participants read the participant information sheet and signed the consent form prior to data collection. The study protocol was approved by the University of Auckland Human Participants Ethics Committee, and conducted in accordance with the Declaration of Helsinki.

### 2.2 Stimuli and design

EEG was recorded during a 30-minute computer task, which entailed the viewing of four types of dynamic image videos: happy face, sad face, neutral face (i.e., mouth opening) and non-biological stimulus (i.e., kaleidoscope). There were four blocks of 40 trials (160 total). In each block, there were 10 happy face, 10 sad face, 10 neutral face and 10 non-biological stimulus videos, presented in random order. Each video was 6000 ms long. Participants were free to rest between the blocks for as long as they wanted.

Happy and sad face videos were taken from the Amsterdam Dynamic Facial Expression Set (ADFES; van der Schalk et al., 2011). The ADFES is freely available for research from the Psychology Research Unit at the University of Amsterdam. Neutral faces were recorded by OK. Past research has validated mouth opening videos of actors as non-emotional (Rayson et al., 2016). Videos used in the present study were made similar to Rayson et al.’s (2016) and the ADFES stimuli in terms of duration, brightness, size, and contrast. Kaleidoscope images presented as the non-biological stimulus were those used in a previous study (Hobson & Bishop, 2016). All stimuli were greyscaled.

Participants were instructed to minimise movement throughout the experiment, and blinking during trials. As *Figure 1* illustrates, each trial started with a 1000 ms fixation cross against a white background. After the fixation cross, the static image stimulus was presented for 2000 ms, followed by a 2000 ms dynamic image in which the expression changed, and ending with a 2000 ms static image of the last frame of the video. Then, a two-alternative forced-choice response slide showing the correct label alongside one of the other three labels prompted the participant to categorise the stimulus as *happy*, *sad*, *neutral* or *other*. The response slide remained on the screen until the participant gave a response using the keyboard. The participant pressed “d” if the label on the left was correct and “k” if the label on the right was correct. In half of the trials, the correct label was on the right, and in half, on the left. Each trial ended with a 1000 ms feedback slide. The feedback slide displayed the word “Correct” or “Incorrect” depending on the key press.

**Figure 1.**
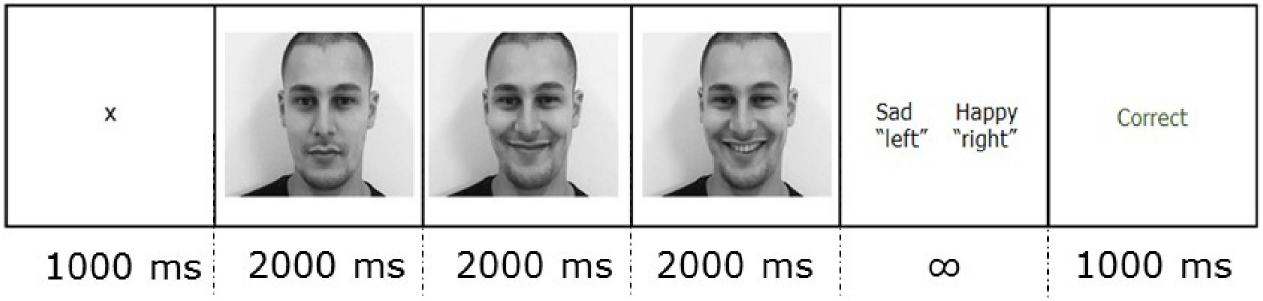
An example of a trial showing the duration of each section in ms. Each condition video was presented for 6000 ms, of which the first 2000 ms was static, the second 2000 ms was dynamic, and the last 2000 ms was static.

Accuracy and reaction time were not analysed. The feedback slide was only used to gauge attention. The highest number of incorrect answers observed for a participant was six (i.e., 4.5%), indicating sustained attention to the stimuli for all participants.

### 2.3 EEG data recording

EEG recording was conducted in an electrically shielded room (IAC Noise Lock Acoustic – Model 1375, Hampshire, United Kingdom) using 128-channel Ag/AgCl electrode nets (Tucker, 1993) from Electrical Geodesics Inc. (Eugene, Oregon, USA). EEG was recorded continuously (1000 Hz sample rate; 0.1-400 Hz analogue bandpass) with Electrical Geodesics Inc. amplifiers (300-MΩ input impedance). Electrode impedances were kept below 40 kΩ, an acceptable level for this system (Ferree et al., 2001). Common vertex (Cz) was used as a reference. Electrolytic gel was applied before the recording started. Each session consisted of two continuous recordings. After the first two blocks, recording was paused and electrolytic gel was re-applied to ensure the impedance was kept low.

### 2.4 EEG data preprocessing and analysis

EEG processing was performed using EEGLAB, an open-source MATLAB toolbox (Delorme & Makeig, 2004). For each participant, the continuous data were downsampled to 250 Hz and then high-passed filtered at 0.1 Hz. 6000 ms conditions starting from the dynamic image onset at time zero were created to get rid of between-session data. Line noise occurring at the harmonics of 50 Hz was removed. Bad channels were identified using the EEGLAB pop_rejchan function (absolute threshold or activity probability limit of 5 SD, based on kurtosis) and interpolated. Data were re-referenced to the average of all electrodes. Infomax ICA was run on each of the preprocessed dataset with EEGLAB default settings. Eye movement and large muscle artefact components were visually identified and rejected for each participant. EEG recordings of three participants were identified as very noisy during the cleaning stage and excluded from further processing.

For each condition, from the 6000 ms image video, 800 ms to 1900 ms early epochs corresponding to the static image and 1950 to 4000 ms late epochs corresponding to the dynamic image were extracted. The analysis was conducted for the mu/alpha band of 8 to 13 Hz over two central clusters of electrodes, six located around C3 on the left hemisphere (i.e., electrodes 30, 31, 37, 41, 42) and six around C4 on the right hemisphere (i.e., electrodes 80, 87, 93, 103, 105), and over six occipital electrodes (O1, Oz and O2). For each of the 15 electrodes, Fast Fourier Transform (FFT) was used to calculate the power spectral density (PSD) in each trial, separately for early and late epochs. For each trial, mu/alpha suppression at each of the 15 electrodes was calculated by taking the ratio of the late epoch PSD relative to the early epoch PSD. Ratio values instead of subtraction values were used as a measure of suppression to control for mu/alpha power variability between individuals that are due to differences in scalp thickness and electrode impedance (Cohen, 2014). Across the central and occipital electrode clusters separately, if a trial had a PSD ratio value greater than 3 scaled median absolute deviations from the median PSD ratio value of the cluster, that trial was excluded as an outlier. For each of the four conditions, the average PSD ratio of the 12 central electrodes was calculated to get a single mu value, and of the three occipital electrodes to get a single alpha value, resulting in eight power scores (i.e., suppression for happy, sad, neutral face and non-biological stimulus images at central and occipital areas) for each participant.

Since ratio data are non-normal, a log transform was used for statistical analysis. A log ratio value of less than zero indicates suppression, zero indicates no change, and greater than zero indicates facilitation.

### 2.5 Source estimation

The 128-channel EEG data were analysed using standardised Low Resolution Electromagnetic Tomography method (sLORETA) source localisation (Pascual-Marqui, 2002; free academic software available at http://www.uzh.ch/keyinst/loreta.htm). sLORETA is an inverse solution that produces images of standardised current density at each of the 6430 cortical voxels (spatial resolution 5 mm) in Montreal Neurological Institute (MNI) space (Pascual-Marqui, 2002). sLORETA images of the mu/alpha band (8-13 Hz) activity during the late epochs of the neutral face and non-biological stimulus conditions were computed for each participant, and then the group averages for the two conditions were extracted. Mu/alpha band power associated with late epochs of the neutral face and non-biological stimulus conditions was compared. A whole-brain analysis was conducted to provide evidence that the reduced mu/alpha band power during the late epoch in the neutral face condition compared to the non-biological stimulus condition was localised to the central instead of posterior regions, indicating stimulus-related differences in sensory and motor activity rather than a cortex-wide activity tapping attention. Voxel-wise *t-*tests were done on the frequency band-wise normalised and log-transformed sLORETA images. For all *t-*tests, the variance of each image was smoothed by combining half the variance at each voxel with half the mean variance for the image. Correction for multiple testing was applied using statistical nonparametric mapping (SnPM) with 5000 permutations.

## 3 Results

### 3.1 EEG results

All statistical analyses were performed using *R studio* (R Studio Team, 2016).

Data from 22 participants were included in the analysis. As explained in the methods section, there were four conditions (i.e., happy face, sad face, neutral face, non-biological stimulus) and two brain regions (i.e., central, occipital) of interest. Before hypothesis testing, a *t*-test for each condition at each brain region was conducted to ensure that the PSD was significantly reduced during the late compared to the early epoch (all *p*-values < .001). Upon confirming suppression in each condition at both brain regions, a 4 × 2 repeated measures ANOVA was conducted. Mauchly’s test indicated that the assumption of sphericity was met for the condition variable. There was a non-significant main effect of condition (*F*(3, 63) = 2.74, *p* = .051, *η*_p_^*2*^ =0.115), and a non-significant main effect of region (*F*(1, 21) = 1.520, *p* = 0.231, *η*_p_*2* = 0.067). However, interpretation of these main effects is qualified by the significant interaction between condition and region (*F*(3,63) = 10.734, *p* < 0.001, *η*_p_^*2*^ =0.338). The interaction effect was investigated further with two sets of pairwise comparisons across conditions at each brain region. Benjamini and Hochberg (1995) false discovery rate (FDR) correction was applied to correct for multiple comparisons between suppression values across the conditions at the central (*p* < .05, FDR corrected) and occipital regions (*p* < .05, FDR corrected). At the central region, only the neutral face condition showed significantly greater suppression than the non-biological stimulus (*p* < .05). Neutral face also showed greater central suppression than the sad face condition (*p* < .05). At the occipital region, suppression was significantly greater in the non-biological stimulus condition than all the other conditions (all *p*-values < 0.05). The distribution of the data points can be seen in *Figure 2*. Three participants had at least one ratio score greater than 1.5 times the interquartile range below the 25th or above the 75th quartile. Removing them did not change the pattern of results, so the analyses are reported including these outliers.

**Figure 2.**
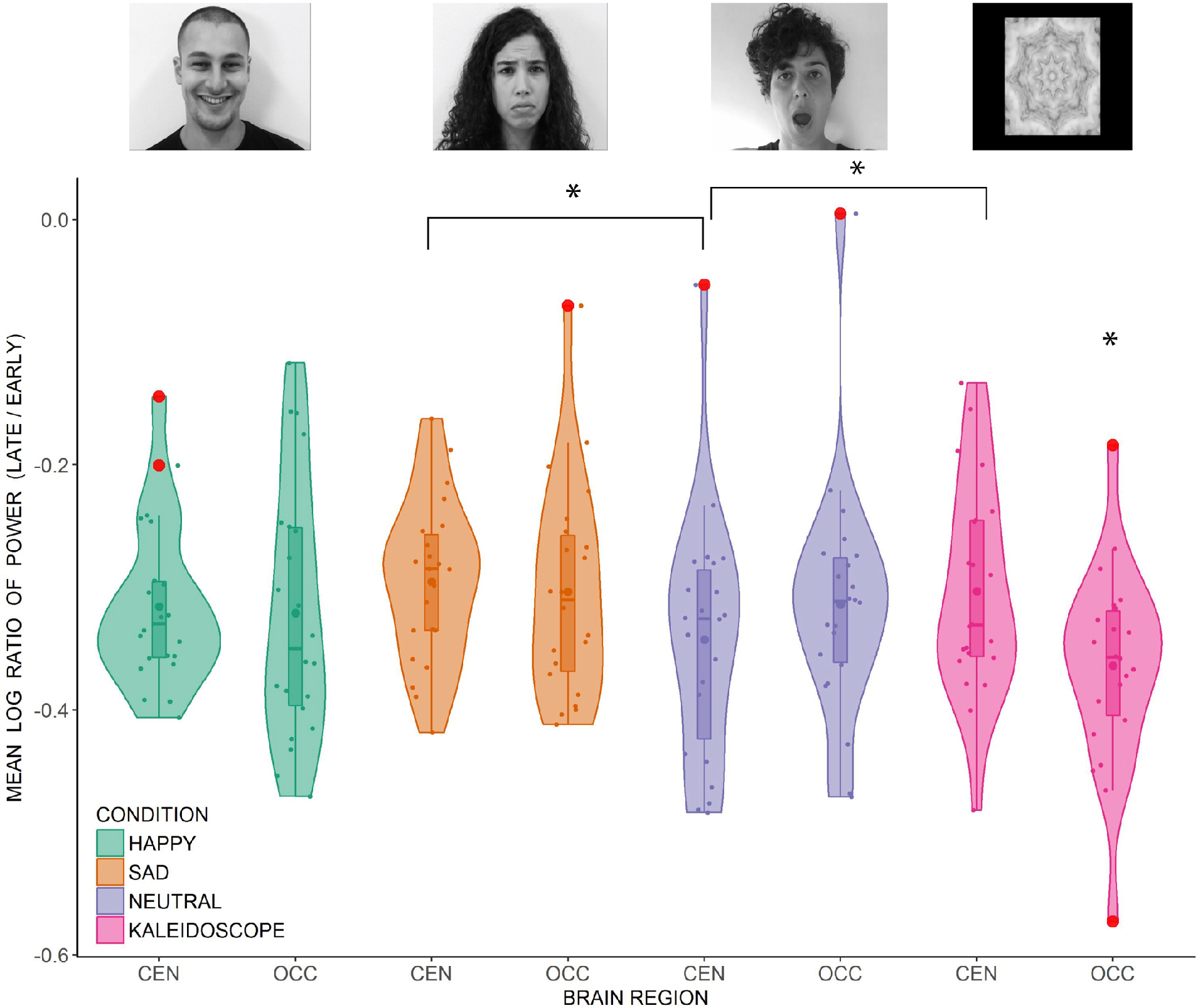
Distribution of the individual mean log ratio scores of 22 participants (dots) in the happy, sad, neutral face and non-biological stimulus conditions at the central and occipital regions. Outliers (>1.5 x interquartile range) are represented by the red disks. Inside the boxplots, dots represent the means and horizontal lines represent the medians. The density plots around the data points represent the kernel probability density of the data at different values. CEN = Central region, OCC = Occipital region. Significant differences are marked by an asterisk (*p* < .05, FDR corrected).

### 3.2 Source Estimation Results

The neural sources of the difference in the mu/alpha band current density power between the neutral and non-biological stimulus conditions during the late epoch (neutral face minus non-biological stimulus) were analysed using sLORETA with a one-tailed test (neutral face < non-biological stimulus). Exceedance proportion test output from sLORETA analysis was used to identify the voxels at which the difference in mu/alpha power between the two conditions was significant (p < 0.05). Based on the exceedance proportion test results which showed a threshold of −3.599 for a p-value of 0.0524, differences in alpha power were localised to the fusiform gyrus (BA20) *t* = −4.03 (X= −55, Y= −40, Z= −30) (MNI coordinates), primary somatosensory cortex (BA3) *t* = −3.80 (X= −40, Y= −25, Z= 40), prefrontal cortex (BA9) *t* = −5.14 (X= 10, Y= 45, Z= 35), and medial premotor cortex (supplementary motor area; BA6) *t* = −3.99 (X= 10, Y= −30, Z= 70) (see *Figure 3*). In the colour scale, blue indicates less alpha power while red indicates the opposite.

**Figure 3.**
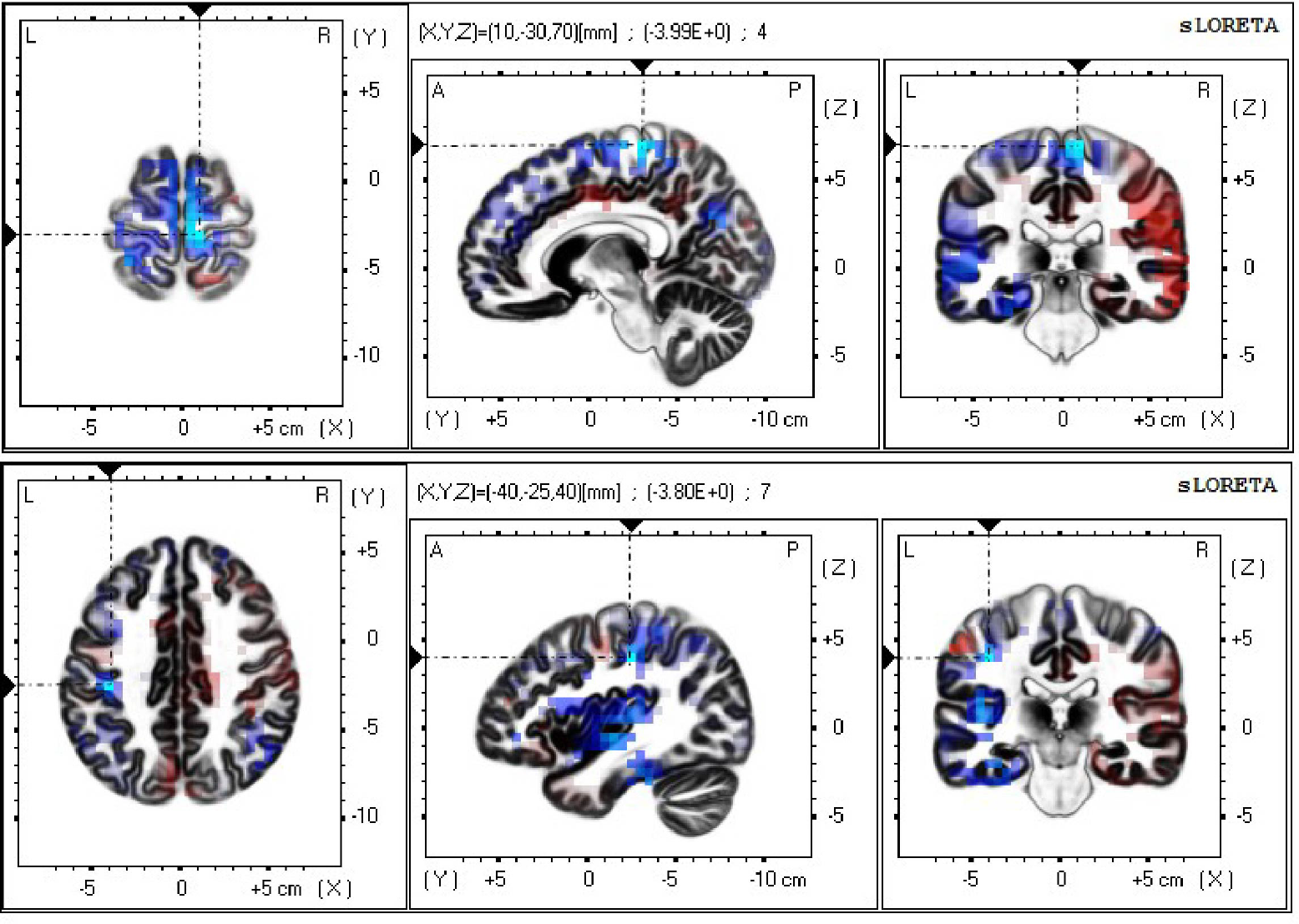
Current density power analysis in the mu/alpha band (8 - 13 Hz), averaged across 22 participants, between the neutral face and non-biological stimulus conditions during the late epoch found significant voxels (*p* < 0.05) best matched to the supplementary motor area (top) and the primary somatosensory area (bottom). Horizontal (left), sagittal (middle), and coronal (right) sections through the voxel with the maximal *t*-statistic (local maximum) are displayed. Blue indicates less power in the alpha band in the neutral face than the non-biological stimulus condition.

## 4 Discussion

The current study investigated the modulation of mu rhythm while participants observed videos of emotional and neutral face movements and non-biological stimulus movements. Mu suppression, but not occipital alpha suppression, was predicted to be greater in the face conditions than the non-biological stimulus condition, with greater suppression in the emotional faces than the neutral face condition. In contrast to our prediction, only the neutral faces were associated with stronger mu activity than that for the non-biological stimulus condition. A lack of difference in mu/alpha band power between the emotional faces and the non-biological stimulus at the central region made it difficult to distinguish mu from posterior alpha modulation during emotional face observation. Greater suppression in the neutral face than the non-biological stimulus condition at the central region accompanied with an opposite pattern at the occipital region suggests that mu rhythm modulation associated with neutral face processing is distinct from the attenuation of the overall alpha activity power associated with information processing and attention. Similar opposing trends of alpha and mu suppression between biological and non-biological movement was observed by Hobson and Bishop (2016). Greater occipital alpha suppression in the non-biological stimulus than the neutral face condition may be explained by low-level visual differences between the two conditions, such as the contrast and the frequency domain information in the stimuli, and/or disparate demands on attention.

In addition to the results from the scalp-recorded EEG activity, source analysis data provide further support for a more localised than an overall difference in the mu/alpha band power between neutral face and non-biological stimulus conditions, suggesting different levels of activity between conditions in the face-related (i.e., fusiform gyrus) and MNN areas, specifically, the primary somatosensory cortex, prefrontal cortex and supplementary motor area. Greater activity in the fusiform gyrus in response to faces than non-biological stimulus was expected as this area responds more to faces than objects (Haxby et al. 2000). The premotor areas, including the supplementary motor area, and the primary somatosensory cortex are the key regions implicated in sensorimotor simulation during action observation (for a review, see Wood et al. 2016) and motor imagery (Burianová et al. 2013; Filgueiras et al., 2018). The premotor cortex has been a primary region investigated in studies of action observation (Buccino et al., 2001; Johnson-Frey et al., 2003; Raos et al., 2004, 2007). While the motor representations of actions are stored in the premotor areas, the somatosensory areas may be involved in storing tactile and proprioceptive representations of these actions (Gazzola & Keysers, 2009). In addition to the role of the somatosensory activity in hand actions (Avikainen et al., 2002; Raos et al., 2004), there is evidence for the involvement of somatosensory representations in our ability to simulate basic emotions while observing facial expressions (Adolphs et al., 2000). Wood et al.’s (2016) review highlights the role of sensory simulation in addition to motor simulation in emotion recognition, pointing to a large overlap between brain areas involved in production and observation of facial expressions. Signalling from the somatosensory cortex to the premotor cortex may be a necessary step for action understanding and imitation (Gazzola & Keysers, 2009). This signalling may explain the significantly less mu/alpha band power present source estimation results show in these two brain areas in the neutral face movement compared to the non-biological stimulus condition.

We offer a number of possible explanations for the EEG results showing the strongest mu suppression to the neutral face movement in the form of mouth opening. Firstly, the results may be attributed to the sensitivity of the sensorimotor cortex to human-object interaction. Most research that has investigated the role of MNN in action observation involves hand and finger movements that almost always suggest some sort of interaction with an object, such as pincer movement with the thumb and the index finger (e.g., Cochin et al., 1999), manipulating objects (e.g., Gazzola & Keysers, 2009), or bringing food to mouth (Ferrari et al., 2003). In addition to limb movements, viewing oro-facial movements has also been observed to induce mu power decrease, with greatest suppression to viewing object-directed actions compared to undirected sucking and biting movements, and least suppression to the viewing of speech-like mouth movements (Muthukumaraswamy et al., 2006). In the present study, the sensorimotor cortex could be engaged by the mouth opening gesture which may have been perceived as an action associated with eating, an action implying interaction with an object (i.e., food), thereby supporting intention understanding (i.e., eating). Secondly, the MNN may be involved in the recognition of deliberate, voluntary gestures rather than involuntary communicative actions. Yet, mu suppression is reported to be modulated by contextual information, such as the actor’s familiarity (Oberman et al., 2008) or their reward value (Gros et al., 2015), or gaming context in which the hand gestures are viewed (Perry et al., 2011). In addition, viewing facial gestures that do not suggest object interaction or deliberate action also seems to modulate mu rhythm (Moore & Franz, 2017; Moore et al., 2012; Rayson et al., 2016; Rayson et al., 2017). Thus, explanations which restrict mu suppression to voluntary or object-related actions are unlikely.

A third explanation is that different types of facial movements may tap different MNN areas. An fMRI experiment conducted by van der Gaag et al. (2007) found bilateral inferior frontal operculum activation to viewing emotional facial expressions but somatosensory activation to neutral movements (i.e., blowing up the cheeks). The authors attributed their findings to distinct processing pathways, more visceral in the former and more proprioceptive in the latter. A similar differential pathway may explain the current findings. Alternatively, if a single mirroring pathway underlies all types of facial movements, greater ambiguity of the action and/or the emotion in the mouth opening image may require the MNN more than the full-blown, easy to recognise emotional expressions. In other words, when the emotion information is presented in high intensity, the cognitive task of recognition may not be demanding enough to activate the MNN, thereby bypassing the whole system, as indexed by the lack of or reduced mu suppression.

Based on recent findings from connectivity research (see, for example, Gardner et al., 2015), our last explanation argues that rather than a global increase/decrease of activity in the totality of the network, a differential modulation of the signalling between the key MNN nodes is more likely to be at work during action observation. There is evidence for the existence of a subgroup of neurons in the human supplementary motor area that is excited by execution, but inhibited by observation of hand grasping actions and facial emotional expressions (Mukamel et al., 2010). These observation-inhibited neurons may be the mechanism for self-other discrimination process related to observing others’ actions, and the strength of their activity may modulate the amount of input from premotor areas to the sensorimotor cortex during action observation (Mukamel et al., 2010; Woodruff et al., 2011). Readily recognisable emotion-related information may activate the observation-inhibited mirror neurons in the premotor areas, leading to less excitatory input to the sensorimotor cortex. On the other hand, neutral facial movements that lack social and emotional information, as in mouth opening, may not activate observation-inhibited neurons as much as easily recognisable expressions do, resulting in stronger excitatory input to the sensorimotor cortex. Thus, in the face of subtle expressions, increased sensorimotor activity may aid action and emotion recognition.

Signalling between and within the key MNN areas during action observation and execution has recently been approached from a Bayesian perspective that suggests the existence of an updating mechanism which continuously attempts to minimise the difference (i.e., the error) between the predicted action and the observed or executed action to achieve an understanding of the most likely cause of an action (Keysers & Perrett, 2004; Kilner et al., 2007b;). According to a predictive coding model of mirror neurons, when the mismatch between the predicted and observed actions of others is large due to the unfamiliarity, unusualness and unexpectedness of the observed action, the network generates a new prediction model, resulting in stronger motor activation (Kilner et al., 2007a, b). In line with this account, several studies have reported greater mu suppression in infants during observation of extraordinary actions (e.g., turning on a lamp with one’s forehead or lifting a cup to the ear) compared to ordinary actions (e.g., turning on a lamp with one’s hand or lifting a cup to the mouth), suggesting that as the deviation of the observed action from the expected action increases, motor activation increases (Langeloh et al., 2018; Stapel et al., 2010). In the current study, the unfamiliarity of the mouth-opening movement as a neutral gesture may have resulted in a greater error signal between the predicted, usual neutral gesture the participants would expect to see, and the observed, unusual neutral gesture they were instructed to categorise as such. Additional predictions that required updating in the mouth opening condition may have activated the sensorimotor areas more than the familiar and ordinary happy and sad gestures. Future research may examine the coordinated activity of the involved brain regions by connectivity analyses to quantify the differences in their associations or dependencies under different conditions.

### 4.1 Limitations

There are several important limitations of the current study that must be noted. Low-level visual properties, such as the contrast and the frequency domain composition of the images, in the face and the non-biological stimulus conditions were not matched. Future studies should aim to match contrast and frequency components of stimuli across conditions in order to mitigate the effect of these non-task related factors on mu/alpha activity. Second, in the face videos, every actor performed only one facial expression. This might have led the participants to learn the movement that followed each static image, leading to habituation across the blocks. Using the same actors for different facial expressions might help avoid habituation-driven mu/alpha activity changes. Another important limitation is related to the uncontrolled degree of movement viewed in each condition. Variability in the amount of movement displayed in videos may have influenced mu and alpha power modulation across conditions. Thus, it is possible that the greater mu suppression to neutral faces reflects more the more pronounced movement in the mouth opening action compared to the happy and sad expressions rather than the differences in social emotional content. Furthermore, face videos used as stimuli may not induce mu modulation that would naturally be observed in real life settings. Finally, the limited sample size and lack of a priori power analysis require further replication studies to shed light on the modulatory influence of observed facial movements on the mu rhythm.

### 4.2 Conclusions

In conclusion, ambiguity or complexity of emotional information may result in greater activity in the sensorimotor areas if difficulty of the emotion recognition task requires a stronger engagement of the simulation system. Present findings provide support for the involvement of the MNN in face simulation, and indicate a complex relationship between sensorimotor activity and facial expression processing. Current data call for further research on the observation-related activity within and between the key brain areas involved in mimicry and social information processing. The explanations offered above which attribute the observed effect to the ambiguity of emotion may be addressed in future studies by comparing the level of activity in the premotor, motor and somatosensory areas in response to social stimuli depicting different intensities of various emotions. High spatial resolution neuroimaging techniques, such as fMRI, can be employed to investigate the involvement of the main MNN areas as well as deeper brain regions in the simulation of ambiguous motor and emotion information.

## Author contributions

OK performed data acquisition, analysis, interpretation of the data and wrote the first draft of the manuscript. IK contributed substantially to the design of the experiment and critical revision of the manuscript. MM contributed substantially to data analysis and critical revision of the manuscript. All authors approved the final draft of the manuscript for publication (OK, MM, IK).

## Acknowledgments

This work was supported by The University of Auckland Postgraduate Research Student Support (PReSS) Account (OK). We thank Veema Lodhia for assistance during data collection and all the participants that took part in the study. This article has been published as a pre-print in bioRxiv (Karakale et al., 2019).

